# Ecological theory applied to environmental metabolomes reveals compositional divergence despite conserved molecular properties

**DOI:** 10.1101/2020.02.12.946459

**Authors:** Robert E. Danczak, Amy E. Goldman, Rosalie K. Chu, Jason G. Toyoda, Vanessa A. Garayburu-Caruso, Nikola Tolić, Emily B. Graham, Joseph W. Morad, Lupita Renteria, Jacqueline R. Wells, Skuyler P. Herzog, Adam S. Ward, James C. Stegen

## Abstract

Stream and river systems transport and process substantial amounts of dissolved organic matter (DOM) from terrestrial and aquatic sources to the ocean, with global biogeochemical implications. However, the underlying mechanisms affecting the spatiotemporal organization of DOM composition are under-investigated. To understand the principles governing DOM composition, we leverage the recently proposed synthesis of metacommunity ecology and metabolomics, termed ‘meta-metabolome ecology.’ Applying this novel approach to a freshwater ecosystem, we demonstrated that despite similar molecular properties across metabolomes, metabolite identity significantly diverged due to environmental filtering. We refer to this phenomenon as ‘thermodynamic redundancy,’ which is analogous to the ecological concept of functional redundancy. We suggest that under thermodynamic redundancy, divergent metabolomes can support equivalent biogeochemical function just as divergent ecological communities can support equivalent ecosystem function. As these analyses are performed in additional ecosystems, potentially generalizable principles, like thermodynamic redundancy, can be revealed and provide insight into DOM dynamics.

## Introduction

Riverine ecosystems receive substantial carbon inputs from terrestrial sources (∼1.9 Pg C yr^-1^), releasing some into the atmosphere and transporting a large portion to the ocean (∼0.95 Pg C yr^-1^)^1,2^. Much of this carbon is dissolved and complexed with other elements as organic matter. As this dissolved organic matter (DOM) travels through watersheds (e.g., along river corridors), it interacts with resident microbial communities and undergoes significant biochemical transformations that influence its fate^1,3–7^. Recent research has suggested that these ongoing biochemical reactions have a significant influence on river corridor biogeochemistry^5,6,8^. Despite the significance of these DOM biochemical reactions, predictive models (e.g., Earth system models, reactive transport models) generally do not represent these detailed processes because they are largely unknown^4,6^. Moreover, the underlying principles governing the detailed chemistry of DOM are under-investigated^5^. Our capacity to predict changes in the functioning of coupled terrestrial-aquatic systems (e.g., watersheds) will be enhanced by resolving these uncertainties^3,7,9^.

Recent studies have continued to elucidate principles governing riverine DOM processing ^5,6,10^. Graham *et al*. 2017^10^ revealed that microorganisms within riverbed sediments preferentially targeted organic molecules based on their thermodynamic favorability, thereby deterministically altering DOM chemistry. Stegen *et al*. 2018^5^ further demonstrated that hyporheic zone metabolism was governed by mixing effects which removed thermodynamic protection (i.e., a “priming effect”). Accordingly, Graham *et al*. 2018^6^ demonstrated that DOM chemistry better predicted microbial respiration rates than community composition, metabolic potential, or expressed metabolisms. Together, these studies indicate a strong connection between DOM chemistry and realized biogeochemical function, and that deterministic processes underlie spatiotemporal variation in DOM chemistry.

The recently proposed synthesis of meta-community ecology and metabolomics, termed “meta-metabolome ecology,” provides new opportunities to deepen understanding of the processes governing DOM chemistry^11^. This framework treats organic molecules in the environment as ‘ecosystem metabolites’ that are both resources for and products of microbial metabolism. A given DOM pool can therefore be thought of as an assemblage of ecosystem metabolites analogous to ecological communities. The framework further suggests that studying the contributions of different ecological assembly processes can offer novel interpretations with biogeochemical implications^11^. To operationalize the conceptual framework, ecological null models can be applied to metabolite assemblages to quantify the relative influences of deterministic and stochastic processes governing metabolome dynamics.

Understanding the relative contributions of deterministic and stochastic processes can help reveal mechanisms driving differences in the molecular properties of DOM pools^11^. Deterministic processes^12^ result from forces in the environment that systematically change the probabilities of observing a given metabolite. This can occur by changing the rates that a given metabolite is produced or transformed, which is analogous to ecological selection changing the birth or death rate of a given biological species. In context of the metabolite assemblages comprising DOM pools, deterministic processes are therefore the outcome of the environment selecting for or against a given metabolite. In contrast, stochastic processes^12^ are the result of random events that lead to uncoordinated increases or decreases in prevalence of individual metabolites. Stochasticity can arise through uncoordinated changes in rates of production or transformation (analogous to random birth/death events in ecological systems) as well as via non-selective transport (analogous to dispersal in ecological systems). Stochasticity dominates when deterministic processes (e.g., selective agents) are not applied consistently through space and/or time, or are too weak to overcome factors such as spatial mixing of metabolites^12–14^.

Further analogies can be drawn to ecological systems whereby stochastic and deterministic processes can be separated into different classes to deepen understanding of the forces governing the molecular properties of metabolite assemblages^11^. As in ecological systems, the influences of deterministic processes can separate into variable and homogenous selection. Variable selection occurs when selective pressures cause assemblages that are separated in space or time to diverge in composition. In turn, differences in metabolite composition are greater than would be expected by random chance^12,13^. In contrast, homogenous selection occurs when selective pressures cause assemblages to have similar composition; differences in metabolite composition are less than expected by random chance^12,13^. A dominant influence of stochastic processes results in differences in metabolite composition that do not deviate from a random expectation^15^. While stochastic processes can also be separated into two classes^12,13^, doing so is beyond the scope of the current study.

Linking the relative influences of variable selection, homogenous selection, and stochastic processes to system dynamics (i.e., hydrology, geochemistry) provides opportunities to better understand spatiotemporal dynamics of metabolite assemblages and inform the representation of DOM chemistry in predictive models. A primary analytical challenge is quantifying the relative influences of deterministic and stochastic processes. As shown in Danczak *et al*.^11^, this challenge can be overcome using metabolite null modeling, which borrows directly from ecological null modeling through the use of dendrograms representing biochemical relationships among metabolites. In ecological systems, null models are often based on phylogenetic and/or functional trait relationships (e.g., Swenson *et al*. 2012^16^, Siefert *et al*. 2013^17^, and Dini-Andreote *et al*. 2015^14^). Using metabolite null modeling, Danczak *et al*.^11^ found that biochemical relationships among metabolites can strongly influence spatial variation in river corridor metabolite assemblages. This points to an opportunity to leverage metabolite null modeling to reveal new principles governing the molecular properties of metabolite assemblages comprising DOM.

Here we use concepts (e.g., stochastic/deterministic processes) and analytical tools (e.g., null models) derived from community ecology to investigate fundamental aspects of metabolite assemblages with respect to (1) within and among assemblage diversity (i.e., alpha and beta diversity), (2) stochastic and deterministic processes governing assemblage composition, and (3) the relationship between stochastic and deterministic processes and metabolite chemistry (e.g., thermodynamic properties and elemental composition). For this, we study the temporal dynamics of both stream and streambed pore water from a low-order river corridor within the HJ Andrews Experimental Forest which has long been the focus of river corridor research^18–21^. This system is representative of steep, low-order river corridors, which dominate headwater river networks both in terms of abundance and relative drainage area^22^ and where riverbed (e.g., hyporheic zone) biogeochemical processes can dominate total respiration^23–25^. We find that despite very similar molecular and thermodynamic properties in bulk DOM pools given by high resolution mass spectrometry (i.e., elemental composition, double-bond equivalent, etc.), deterministic processes drove divergence in the biochemical transformations connecting metabolites, both between and within surface and pore waters. Furthermore, our results point to a new concept referred to as ‘thermodynamic redundancy’ in which spatially or temporally separate metabolite assemblages have indistinguishable thermodynamic properties despite divergence in other metabolome characteristics.

## Results

### Metabolite properties were similar across surface and pore water

Given that the sampled surface water had likely passed through the subsurface multiple times within the studied field system^21,26,27^, we expected metabolite assemblages within the surface and pore water to share some molecular properties. This was borne out with respect to properties inferred directly from assigned molecular formulae. More specifically, the surface and pore water metabolite assemblages had similar thermodynamic and molecular properties (**Figure 2**). The standard Gibb’s Free Energy of carbon oxidation (ΔG°_cox_), double-bond equivalents (DBE), and modified aromaticity index (AI_Mod_) did not significantly differ between surface and pore water (p-value > 0.05). While the thermodynamic and molecular properties varied through time, they did not clearly follow diel hydrological dynamics (**Figure 1**). Similarities in thermodynamic and molecular properties between surface and pore water may be due to significant hydrologic connectivity in the study system^19–21^. This mixing has the potential to minimize the signatures of organic matter processing within surface or subsurface domains. Follow-on analyses reveal that mixing does not, however, fully overcome the signatures of localized processes (as discussed below).

**Figure 1:**
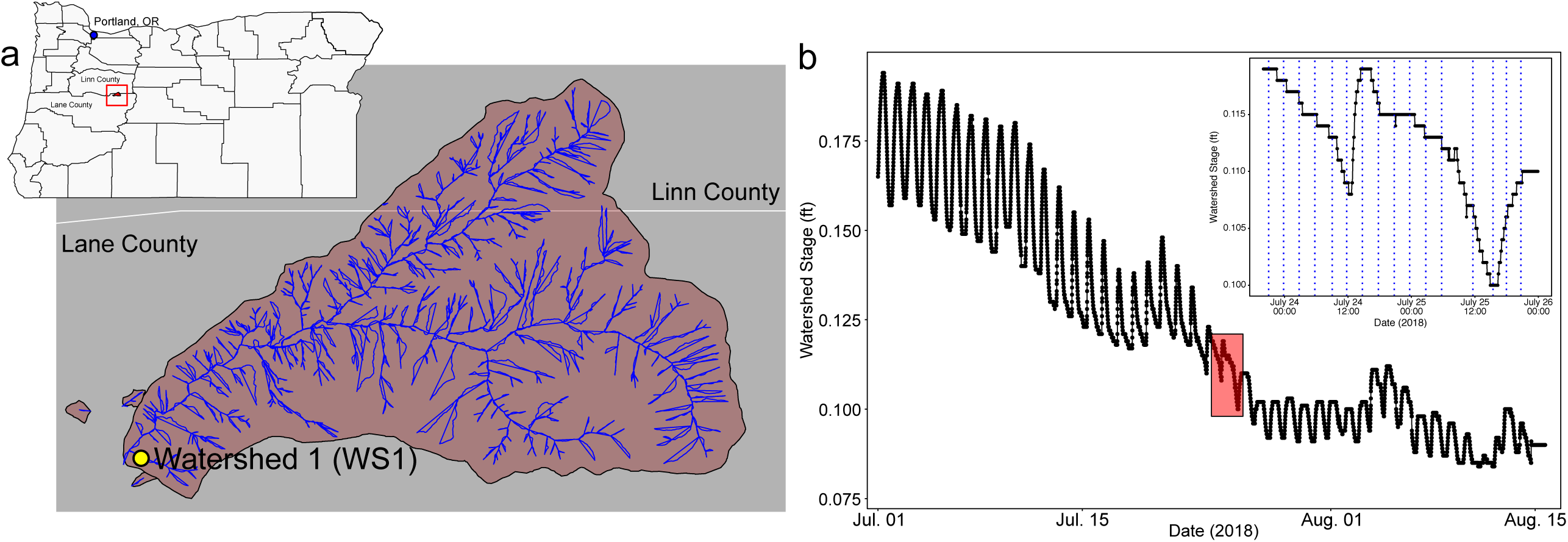
An outline of the study site and associated hydrology. a) Map of Watershed 1 within the HJ Andrews Experimental Forest in Oregon. b) Hydrograph for Watershed 1 with the sampling period highlighted in red and expanded upon in the inset. Sampling points are indicated by the blue dashed lines in the inset.

**Figure 2:**
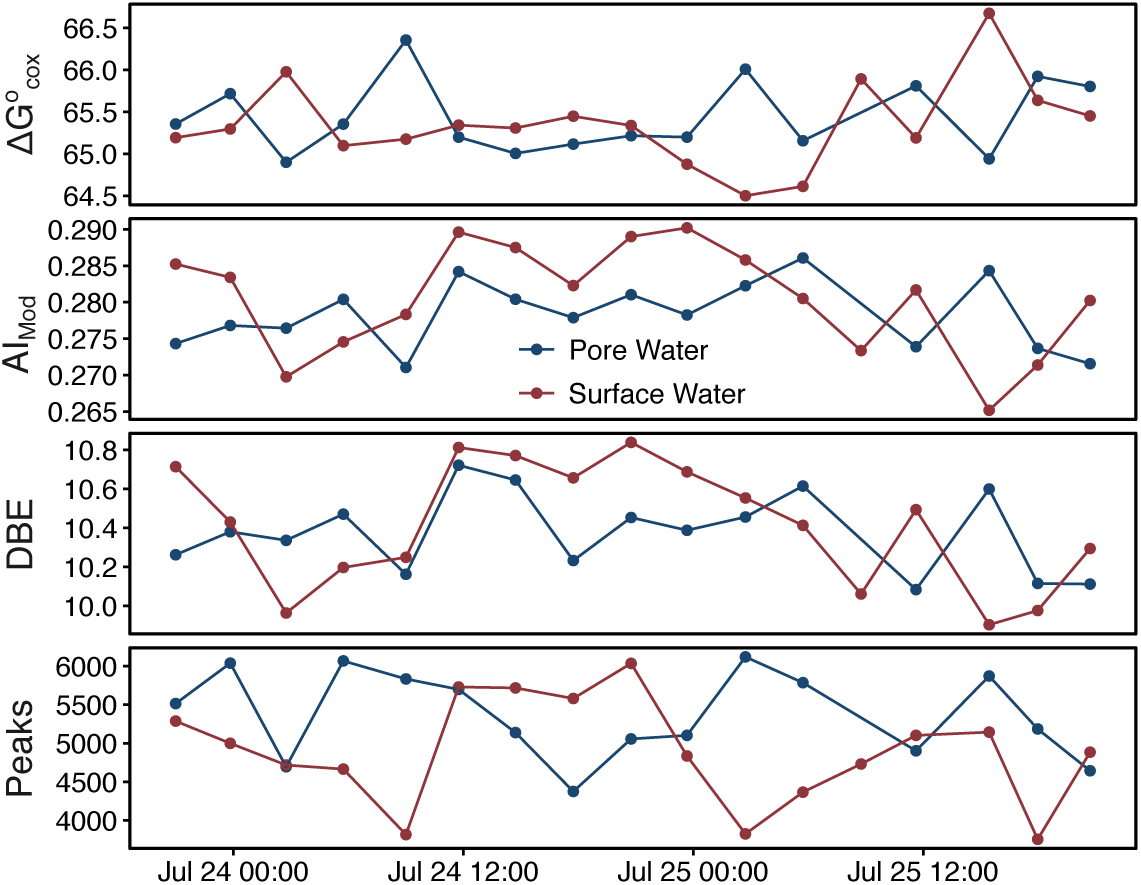
Plots of average chemical properties through time, separated by environment (i.e., pore water and surface water). Abbreviations are as follows - standard Gibb’s Free Energy of carbon oxidation (ΔG°_cox_), modified aromaticity index (AI_Mod_), and double-bond equivalents (DBE). Peak counts refer to the number of peaks within a given sample.

### Conserved alpha diversity and molecular properties contrast with divergence in composition, revealing thermodynamic redundancy

Additional analyses examining both within metabolome diversity (i.e., alpha diversity) and among metabolome differences in composition (i.e., beta diversity) presented an apparent contradiction; metabolomes with similar within-metabolome properties and diversity had divergent composition. This leads to the proposed concept of thermodynamic redundancy, discussed below. More specifically, the dendrogram-based alpha diversity values were largely similar between surface and subsurface metabolomes mirroring dynamics in molecular and thermodynamic properties (**Figure 3**). Patterns of Faith’s PD mostly followed molecular property patterns, indicating that there were no major differences in dendrogram structure between surface and pore water metabolomes (p-value: 0.063). Other alpha diversity metrics that use dendrogram-based relational information (i.e., MPD, MNTD, VNTD, VPD) followed similar trends between surface and pore water metabolomes (p-value: > 0.1). These results indicate that across surface and porewater there are conserved molecular properties and biochemical transformation network topologies, both of which are used to estimate the dendrogram used for alpha diversity analyses. Alpha diversity analyses do not, however, directly evaluate variation in composition across metabolomes. Beta-diversity metrics can be used to make such comparisons.

**Figure 3:**
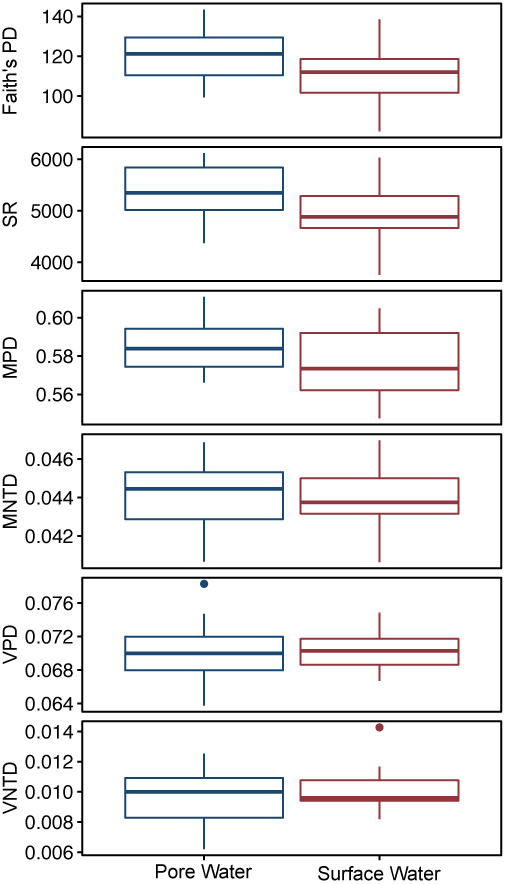
Boxplots illustrating metabolome alpha diversity. Abbreviations are as follows – Faith’s Phylogenetic Diversity (PD), species richness (SR), mean pairwise distance (MPD), mean nearest taxon distance (MNTD), variation of pairwise distance (VPD), and variation in nearest taxonomic distance (VNTD). If a p-value is listed, significant differences were identified using a Mann Whitney U test.

Comparison of metabolome assemblages using beta diversity metrics revealed significant divergence in metabolome composition, despite the high degree of similarity in alpha diversity. More specifically, Jaccard dissimilarity and β-mean nearest taxon distance (βMNTD) principal coordinate analysis (PCoA) plots showed clear separation between surface and subsurface metabolomes (**Figure 4**; Jaccard p-value – 0.005; βMNTD p-value – 0.02). Furthermore, the Jaccard-based analyses reveal significantly greater differences than did βMNTD. This reflects patterns observed within the dendrograms used in the estimation of βMNTD, but not used to estimate Jaccard; similarities in molecular properties were captured in the dendrogram resulting in decreased separation across the βMNTD ordination, relative to the Jaccard-based PCoA. Taken together, these results demonstrate that metabolite profiles with indistinguishable molecular and thermodynamic characteristics, as well as similar levels of alpha diversity, can nonetheless be composed of different metabolites when viewed at the level of specific metabolites. This opens the possibility that localized—and potentially temporally variable—deterministic processes drive spatiotemporal variation in metabolite assemblages, which ultimately result in habitat-specific metabolomes.

**Figure 4:**
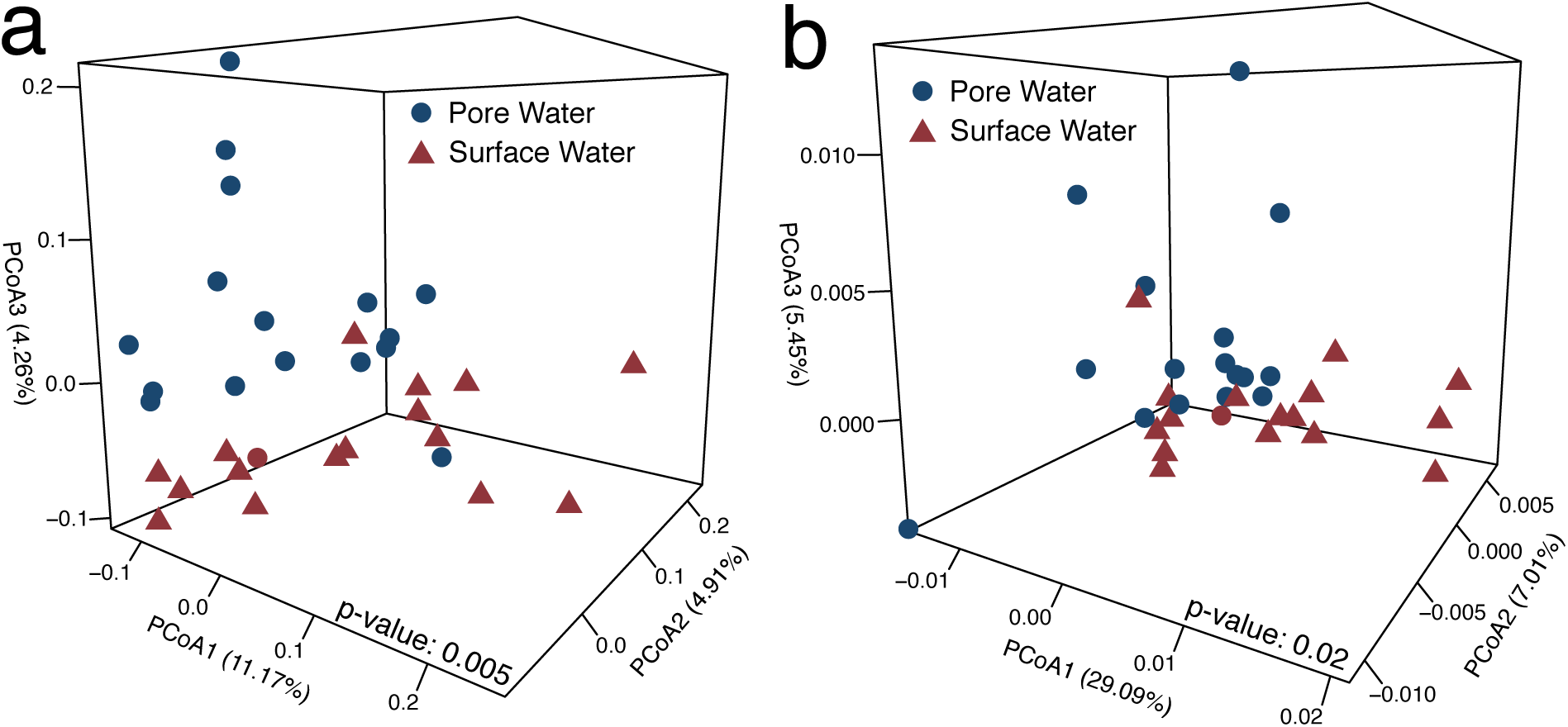
Metabolome beta diversity principal coordinate analyses (PCoA). a) Jaccard dissimilarity-based PCoA b) βMNTD-based PCoA. Significant differences between groups (i.e., pore water and surface water) were determined using PERMANOVA and are indicated in the bottom graph.

Results discussed above present an apparent contradiction whereby there is divergence in composition, but consistency in thermodynamic/molecular properties and alpha-diversity metrics. To reconcile these outcomes, we propose the concept of thermodynamic redundancy. Conceptually, thermodynamic redundancy is similar to the ecological observation of functional redundancy, whereby different biological taxa can fill the same functional role. In the case of thermodynamic redundancy, different metabolite assemblages are comprised of different metabolites (analogous to biological taxa) but have similar thermodynamic and molecular properties. Given strong influences of DOM thermodynamics in river corridors, we propose the hypothesis that thermodynamic properties of individual metabolites are analogous to functional roles of biological taxa.

From an ecological perspective, functional redundancy has been observed repeatedly in both microbial communities (with respect to metagenomic profiles) and macro-organisms such as plant communities (with respect to functional traits such as specific leaf area)^28–31^. For example, the human gut can have numerous different steady state microbial communities that all exhibit healthy function due to redundant metabolisms^32^. We hypothesize that this analogy extends to metabolites in that different assemblages may support the same biogeochemical function (e.g., net rate of denitrification) by meeting some given thermodynamic requirements. Alternatively, thermodynamic redundancy may instead capture the biogeochemically-relevant historical processes that led to metabolomes with similar molecular properties but divergent composition, rather than true functional diversity.

The degree to which thermodynamic redundancy is observed across metabolite assemblages will require data from a broad suite of environmental systems. It will be important to evaluate this concept with paired measured biogeochemical rates and with more detailed metabolome data that include information on molecular structure to assess its impact on the potential functional role of organic metabolites. Regardless of the degree to which thermodynamic redundancy indicates true functional redundancy, extending the general concept of redundancy to metabolomes further emphasizes the significant breadth of conceptual parallels between ecological communities and metabolite assemblages.

### Divergence in metabolite assemblages was associated with biochemical transformations

The concept of thermodynamic redundancy indicates conserved thermodynamic properties despite strong divergence in metabolite composition. Through additional multivariate analyses we found that this divergence was driven by transformations that were used to define biochemical relationships among metabolites in our analyses. More specifically, through a Jaccard dissimilarity-based NMDS analysis we found that profiles of biochemical transformations were divergent between surface and subsurface metabolomes (**Figure 5**; p-value: 0.0082). Examining the transformations by elemental composition showed that transformations containing only C, H, and O were significantly more frequent within surface water (p-value: 0.014) while N-containin transformations (including the loss or gain of amino acids) occurred more frequently within the pore water (p-value: 0.008). Previous work has also shown greater abundance of N-based transformations in pore water, relative to surface water^5^. While we can only speculate, these results suggest that generalizable principles might exist in terms of how biochemical transformations vary between surface and pore water.

**Figure 5:**
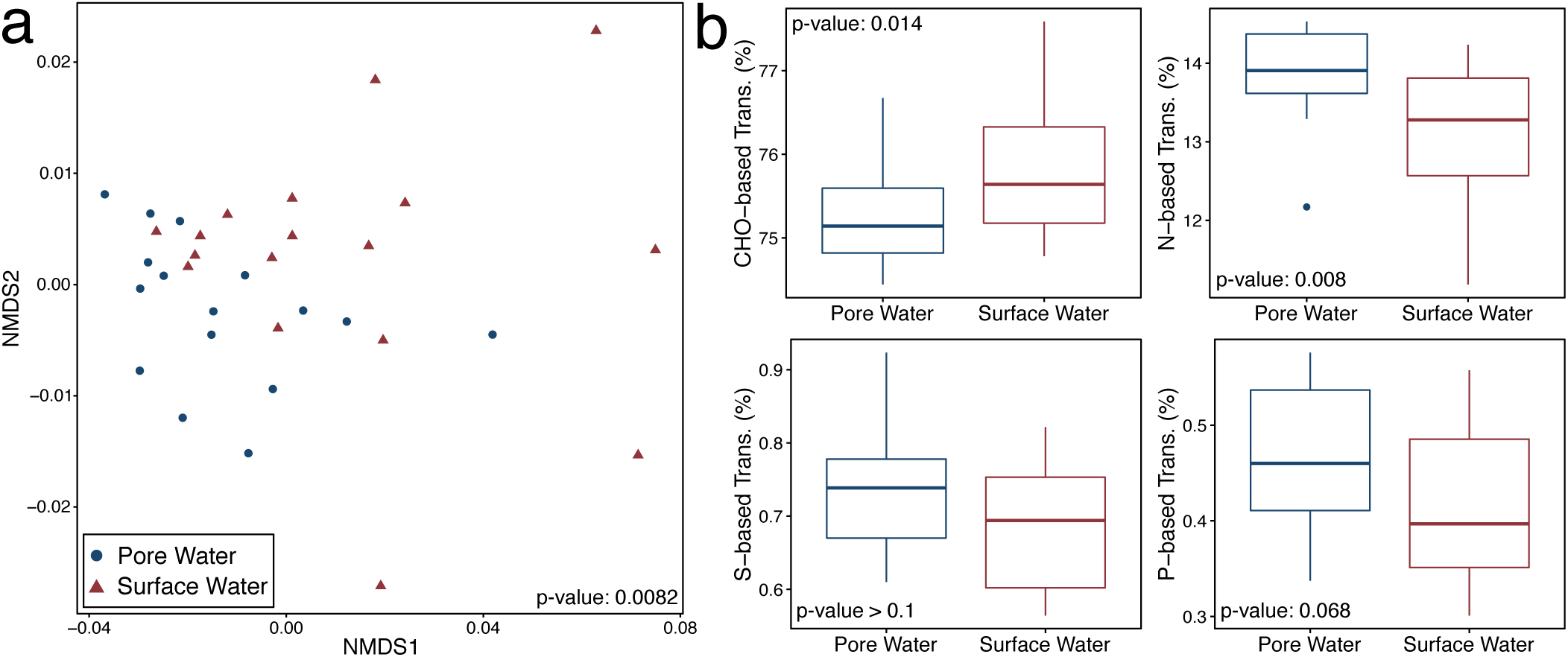
An investigation of potential biochemical transformations throughout the watershed. a) Jaccard-based non-metric multidimensional scaling (NMDS) graph for transformation profiles, with significant differences between groups determined by PERMANOVA and indicated in the bottom right. b) Boxplots comparing the relative proportion of transformations with specific elemental compositions observed within pore or surface water. Significance indicated by Mann Whitney U tests are indicated in the bottom or top left. For example, the surface water had a significantly higher proportion of transformations containing only C, H, and O than the pore water.

As in Stegen *et al*.^5^, we suggest that the subsurface has a greater capacity for biomass turnover and proteolytic activity due to increased microbial load as compared to the surface water. We also suggest that the higher frequency of N-transformations in the pore water were not due to differences in N limitation causing enhanced N mining given that N concentrations (e.g., NO_3_, NO_2_, and total N) were below our limit for detection in both surface and pore water (**Supplemental File 1**). However, we did not measure organic N, so we cannot exclude the possibility that N was more limited in the subsurface than the surface due to potentially greater microbial load. Alternatively, these differences could arise from hotspot activity which has been reported within other riverbed sediments/hyporheic zones^6^. Regardless of the mechanism, the consistency between this study and previous work suggests that shallow subsurface domains (often associated with hyporheic zones) may consistently be characterized by greater abundance of N-containing biogeochemical transformations. Multi-system comparative studies will be needed to evaluate this possibility, which could emerge as a principle that is transferable across river corridor systems, providing an opportunity to inform the structure of mechanistic predictive models.

### Deterministic processes drove differences between surface and pore water metabolite assemblages

Divergence in metabolite assemblage composition through space or time can be due to stochastic processes, deterministic processes, or some combination of the two. Deterministic processes can have strong influences when biotic or abiotic features cause systematic differences in organismal reproductive success or metabolite expression across assemblages^15^. Stochastic processes can arise due to spatiotemporal differences whereby random or uncoordinated ‘demographic events’ (i.e., organismal birth/death or metabolite expression/transformation) lead to divergence in composition that is not due to systematically imposed deterministic factors^13,33^. Stochastic processes can also be dominant when there is significant movement or mixing of organisms/metabolites across spatial locations (i.e., across ecological communities or metabolite assemblages). The β-nearest taxon index (βNTI) metric, a phylogenetic null modeling approach, has been shown to quantitatively estimate the relative contributions of these stochastic and deterministic processes^12,13,15^. This provides much deeper insight into the mechanisms driving observed spatiotemporal patterns in community/assemblage composition when compared to more traditional methods such as ordinations, redundancy analysis, or regressions.

Applying null modeling approaches to metabolite assemblages showed that divergences observed through ordination analysis (**Figure 3**) were overwhelmingly due to deterministic processes that arise from differences in abiotic and/or biotic features. Specifically, the deterministic processes observed here were akin to the concept of ‘variable selection’ in ecological communities. Variable selection can dominate the assembly of communities when features of the environment systematically drive divergence in composition by causing spatial or temporal shifts in the relative fitness of different biological taxa. We infer that an analog to variable selection driven by features in the biotic and/or abiotic environment is causing divergence in metabolite assemblages within our study system despite conserved levels of alpha-diversity and molecular properties (**Figures 2 and 3**). It is important to recognize that this is not a pre-determined outcome of sampling different locations within the river corridor. The divergence between surface and porewater metabolite assemblages could have been due to limited exchange enabling compositional divergence to arise through uncoordinated (i.e., stochastic) changes in metabolite production and transformation. Such a scenario would have been akin to dispersal limitation enabling ecological drift, which is itself akin to genetic drift within the theory of population genetics^34^. Recent application of the βNTI null model to river corridor metabolite assemblages from the mainstem of the Columbia River showed that such stochastic scenarios are possible and potentially likely^11^.

Examining dynamics within surface or porewater revealed stronger influences of deterministic processes in porewater (relative to surface water), suggesting highly localized biotic or abiotic processes with very strong influences over assemblage composition. Furthermore, porewater metabolomes were more consistently governed by variable selection than those in surface water (**Figure 6**; p-value: < 0.001). This was true despite the study system appearing to be well-mixed, whereby advective transport of water-soluble metabolites could overwhelm deterministic processes causing compositional divergence (akin to ‘mass effects’ in ecological meta-communities)^13,35^. Based on correlations with other physical and chemical variables, deterministic pressures within the surface water seem to be associated with geochemical conditions, including sulfate and dissolved oxygen concentrations (**Supplemental File 2**). No physical or chemical variables were significantly related to the level of determinism associated with porewater metabolite assemblages. These results suggest that different biogeochemical processes are at play in surface and subsurface domains, despite the surface water being an integration of pore water through space and time^18–21^.

**Figure 6:**
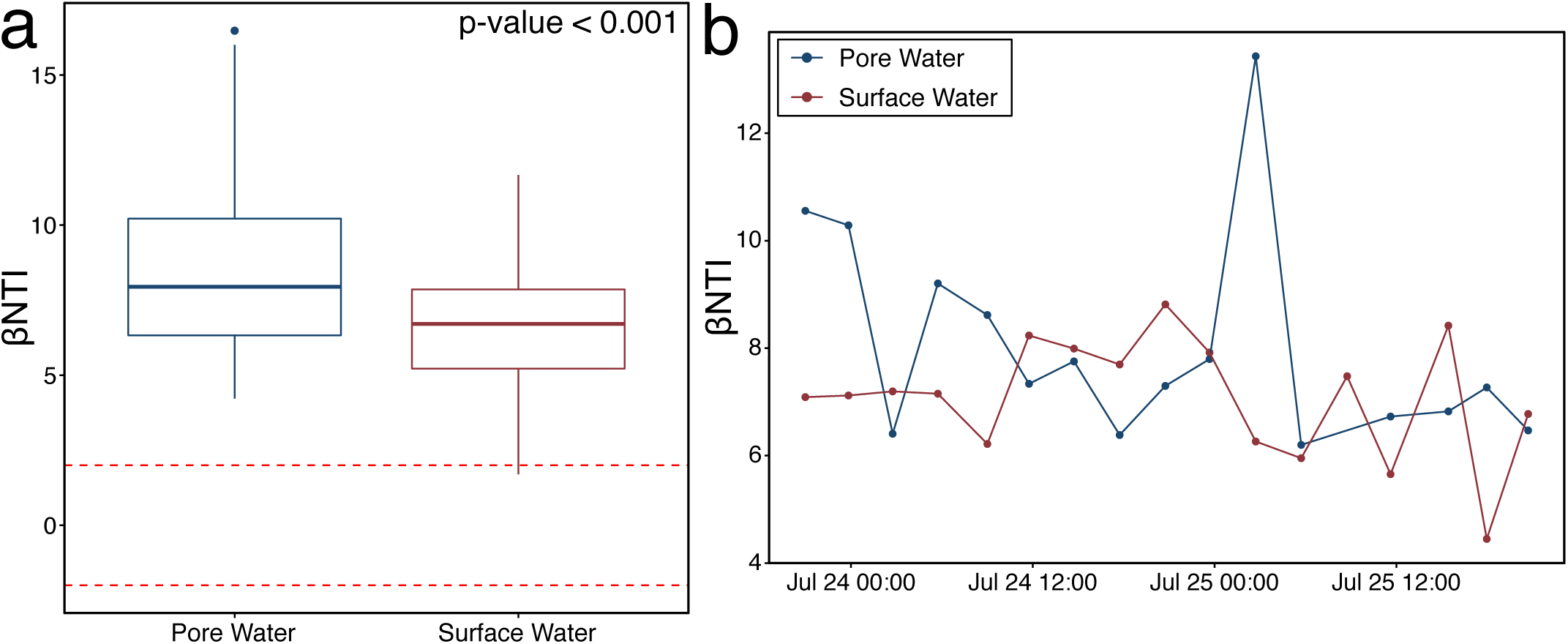
β-nearest taxon index (βNTI) calculations across the watershed. a) Boxplots illustrating differences in βNTI results. Mann Whitney U test significance is indicated in the upper right corner. b) Mean βNTI for each time point separated by water type.

One of the key biogeochemical differences between surface and subsurface domains in the study system and in other river corridors^5^ is the variation of putative biochemical transformations. This inference is supported through analyses linking these putative biochemical transformations to influences of deterministic processes. The relative frequencies of many individual biochemical transformations, regardless of the molecule gained or lost, were significantly correlated to the level of determinism. For most transformations, these correlations were similar between surface and pore water metabolite assemblages (**Supplemental File 3**). Grouping transformations by elemental compositions as above, however, revealed that determinism in the surface water was positively associated with N-, S-, and P-containing transformations and negatively related to those transformations containing only C, H, and O. These results indicate that as N-, S-, and P-containing transformations become more frequent within the surface water, overall metabolome composition begins to diverge. Within the porewater, only S-containing transformations were significantly positively related to deterministic processes. The absence of a strong N-containing transformation correlation within the porewater contrasts with the overall frequency dynamics discussed earlier and likely points to more complex organic N metabolism. To further reveal underlying processes and their dynamics will require more detailed geochemical (e.g., dynamics of vertical redox gradients) and molecular investigations (e.g., metatranscriptomics of microbial communities), likely across other river corridors and longer time periods.

## Discussion

A key element limiting accurate representation of DOM cycling in predictive models is understanding the processes governing spatiotemporal variation in metabolite assemblages and the follow-on impacts to emergent biogeochemical function. To address this challenge, we took a novel approach based on concepts and methods from metacommunity ecology. We find that deterministic processes drive divergence in metabolite assemblage composition through both space and time. This divergence was observed despite similar alpha diversity and molecular/thermodynamic properties. We also provide evidence that deterministic processes which cause metabolome divergence are associated with organic transformations. This indicates that expressed microbial metabolisms should be highly dynamic in time and should diverge between surface and subsurface components of the river corridor. Given strong similarity in molecular properties across surface and subsurface domains, we further propose that divergent metabolite assemblages have the potential to be thermodynamically equivalent.

This highlights a major, unresolved question that is fundamental to understanding the role of environmental metabolites—as both reactants and products—in emergent biogeochemical function. That is, what are the processes that give rise to observed metabolite assemblages and what is the interplay of these processes with biogeochemical function? Future work should focus on understanding the degree to which variation in the composition of metabolite assemblages influences variation in biogeochemistry irrespective of changes in molecular properties. This is analogous to the question of how important microbial community composition is to realized biogeochemical function^6,36,37^. It is often found that microbial composition itself is not a primary driver of biogeochemical function, which indicates a significant amount of functional redundancy^6,38,39^. In other cases, however, microbial community composition corresponds well with ongoing biogeochemical processes. For example, arsenic mobilization within contaminated soils in Bangladesh was driven by the presence and distribution of diverse taxa associated with arsenic and iron reducing bacteria^40^.

Similar functional profiles despite divergent taxonomic composition is a common feature in ecological systems^31,37,41^. Different microbial communities within the human gut or in soil environments will provide similar (if not indistinguishable) contributions to overall ecosystem function^32^. Analogously, divergent metabolite assemblages can have indistinguishable thermodynamic and molecular properties, though this does not necessarily indicate that the metabolite assemblages are identical with respect to biogeochemical function. Both the surface and pore water metabolite assemblages had conserved thermodynamic and molecular properties but were compositionally divergent due to strong deterministic processes (**Figure 6**). This suggests that compositionally divergent metabolite assemblages could be redundant with respect to bulk biogeochemical processes (e.g., respiration rates) that have been shown to be influenced by metabolite thermodynamics^6,42^. Whether these outcomes are driven by differential substrate preference across the riverbed or common labile carbon depletion, the divergence in metabolite assemblages suggests that these environments can take different paths while maintaining similar bulk chemical and thermodynamic properties. In other words, redundancy appears to exist at higher levels of metabolite properties, but not at the lower levels associated with biochemical linkages among metabolites. An open question is the degree to which net biogeochemical rates respond to higher-level properties (e.g., thermodynamics of individual metabolites) versus lower-level biochemical mechanisms. Evaluating this question is fundamental to understanding whether and how thermodynamic redundancy is association with redundancy of biogeochemical function.

Metabolite assemblages are examined as snapshots of the organic compounds at a given point in time and space. By analyzing assemblages together and viewing them as analogs to ecological communities, we can draw upon the concepts, theory, and tools developed with meta-community ecology. Doing so provides novel insight into the processes that shape spatiotemporal dynamics of metabolite assemblages. Here, using this approach we found that variable selection can dominate spatial and temporal dynamics of metabolite assemblages, potentially via underlying biochemical processes associated with dynamic organic N, S, and P metabolism. Similarities between this study and previous work hint at the potential to elucidate generalizable principles that could be used to enhance the predictive capacity of process-based simulation models (e.g., reactive transport codes). Applying our analytical framework to ecosystem metabolomes from a broad suite of river corridors and pairing these analyses with biogeochemical rate measurements will provide exciting opportunities to test and reveal generalizable principles.

## Methods

### Site Description

Samples for this study were collected from Watershed 1 (WS01) in the HJ Andrews Experimental Forest, Oregon, USA (**Figure 1**)^19,21^. For a detailed description of this study site, please refer to Ward *et al*.^21^ and Wondzell *et al*.^19^. Briefly, WS01 is a shallow, low-order, headwater stream which is hydrologically connected to the surrounding terrestrial environment^19,21^. The river corridor is forested, and evapo-transpiration drives diel fluctuations in stream discharge (**Figure 1**)^26,27^. Given that these hydrologic dynamics occur with regular frequency, they offer an opportunity to study changes in DOM composition through time in both the surface water and pore water. This study was conducted under low-discharge conditions during July 23-25, 2018, when diel stage fluctuations can cause spatially intermittent flows, the proportion of total down valley flow passing through the hyporheic zone is maximized, and connectivity between the subsurface and surface was the highest. Therefore, the surface water collected at the sampling location has likely passed through the hyporheic zone multiple times^21,26,27^.

### Sample Collection

Three points separated by ∼4 meters were selected along the river corridor to collect pore water samples. Approximately 20 mL of pore water was collected from each of these locations every 3 hours for 48 hours. Concurrently, surface water was collected in triplicate from the same spatial position as the central pore water location. In total, 102 total samples were collected over 17 time points. Surface water was collected using a 60 mL syringe through Teflon tubing while the pore water was collected using a syringe attached via Teflon tubing to a 30 cm long stainless-steel sampling tube (MHE Products, MI, USA) with a slotted screen across the bottom ∼5cm. One sampling tube was installed to 30cm depth at each pore water sampling location; these tubes remained in place during the 48-hour time course of sampling. Prior to sampling a given location, the syringe was flushed 3 times with the source water to ensure only the desired water was collected. All samples were filtered through a 0.2 μm Sterivex filter (Millipore, MA, USA). At each time point, one filter was used for all pore water samples, and a different filter was used for the surface water. To minimize contamination, water passing through a given filter was collected for analysis using a needle attached to the filter and injected through a septum. During sampling, water temperature, approximate water stage, and pH were measured. Water samples for DOM analysis were injected into amber borosilicate glass vials. Samples for cations and anions were injected into clear borosilicate glass vials. Once collected, samples were stored in a cooler on blue ice until they could be frozen until they were processed in the lab.

### Geochemistry

Anion concentrations were measured using a Dionex ICS-2000 anion chromatograph with AS40 autosampler using an isocratic method (guard column: IonPac AG18 guard, 4×50mm; analytical column: IonPac AS18, 4×250mm; suppressor: RFIC ASRS, 300 rmm, self-regenerating; suppressor current: 57mA). The isocratic method was a 15-minute run with a 1 mL/min flow rate with 22 mM KOH at 30 degrees C and 25 μL injection volume. Standards were made from Spex CertiPrep (Metuchen, NJ, 08840) 1000 mg/L anion standards. NO2 standard was diluted in the range of 0.04 to 20 ppm. F standard was diluted in the range of 0.2 to 10 ppm. Cl and SO4 standards were diluted in the range of 0.16 to 80 ppm. NO3 standard was diluted in the range of 0.12 to 60 ppm. Ion peaks were identified and integrated manually in the software.

Cations samples were prepared with nitric acid. Samples were measured with a Perkin Elmer Optima 2100 DV ICP-OES with an AS93 auto sampler. A Helix Tracey 4300 DV spray chamber and SeaSpray nebulizer were used with double distilled 2 % nitric acid (GFS Chemicals, Inc. Cat. 621) and a flow rate of 1.5 mL/min. Calibration standards were made with Ultra Scientific ICP standards (Kingstown, RI). P, Mg, Ca, K, and Na standards were diluted in the range of 5-4000 ppm. Fe standard was diluted in the range of 0.5-400 ppm.

Non-purgeable organic carbon (NPOC) was determined by a Shimadzu combustion carbon analyzer TOC-L CSH/CSN E100V with ASI-L auto sampler. An aliquot of sample was acidified with 15% by volume of 2N ultra-pure HCL. The acidified sample was then sparged with carrier gas for 5 minutes to remove the inorganic carbon component. The sparged sample was injected into the TOC-L furnace at 680°C using 100 uL injection volumes. The best 3 out of 4 injections replicates were averaged to get final result. The NPOC organic carbon standard was made from potassium hydrogen phthalate (Nacalia Tesque, lot M7M4380). The calibration range was 0.5 to 10 ppm NPOC as C.

### Fourier Transform Ion Cyclotron Resonance Mass Spectrometry Sample Preparation, Data Collection, and Data Preprocessing

Fourier Transform Ion Cyclotron Resonance Mass Spectrometry (FTICR-MS) was used to provide ultrahigh resolution characterization of metabolite assemblages within each DOM sample. Aqueous samples (NPOC 0.33-0.99 mg C/L) were acidified to pH 2 with 85% phosphoric acid and extracted with PPL cartridges (Bond Elut), following Dittmar *et al*.^43^. Subsequently, high-resolution mass spectra of the organic matter were collected using a 12 Tesla (12T) Bruker SolariX Fourier transform ion cyclotron resonance mass spectrometer (Bruker, SolariX, Billerica, MA) located at the Environmental Molecular Sciences Laboratory in Richland, WA. Samples were directly injected into the instrument using a custom automated direct infusion cart that performed two offline blanks between each sample. The FTICR-MS was outfitted with a standard electrospray ionization (ESI) source, and data was acquired in negative mode with the needle voltage set to +4.4kV, resolution was 220K at 481.185 m/z. Data were collected with an ion accumulation time of 0.08 sec and 0.1 sec from 100 m/z – 900 m/z at 4M. One hundred forty-four scans were co-added for each sample and internally calibrated using OM homologous series separated by 14 Da (–CH2 groups). The mass measurement accuracy was typically within 1 ppm for singly charged ions across a broad m/z range (100 m/z - 900 m/z). BrukerDaltonik Data Analysis (version 4.2) was used to convert raw spectra to a list of m/z values by applying the FTMS peak picker module with a signal-to-noise ratio (S/N) threshold set to 7 and absolute intensity threshold to the default value of 100. Chemical formulae were assigned using Formularity^44^, an in-house software, following the Compound Identification Algorithm ^45–47^. Chemical formulae were assigned based on the following criteria: S/N >7, and mass measurement error < 0.5 ppm, taking into consideration the presence of C, H, O, N, S and P and excluding other elements. This in-house software was also used to align peaks with a 0.5 ppm threshold.

The R package ftmsRanalysis^48^ was then used to remove peaks that either were outside the desired m/z range (200 m/z – 900 m/z) or had an isotopic signature, calculate a number of derived statistics (Kendrick defect, double-bond equivalent, aromaticity index, nominal oxidation state of carbon, standard Gibb’s Free Energy of carbon oxidation), and organize the data into a common framework ^49–52^. Samples that were run at both ion accumulation times were combined; given that different IATs will detect different compounds^53^, by combining the two IATs we can gain a more complete characterization of the metabolite assemblages. Replicates were further combined such that if a metabolite was present in one replicate, it was included in the composite assemblage. Because peak intensities cannot be used to infer concentration, all peak intensities were changed to binary presence/absence. In turn, observing a metabolite in multiple replicates was equivalent to observing it in a single replicate; the absence of a peak is defined as below the limit of detection. One sample (PP48_000012) was considered an outlier

### Metabolite Dendrogram Estimation

A transformation-weighted characteristics dendrogram (TWCD) was generated following the protocol outlined in Danczak *et al*.^11^. First, biochemical transformations were identified within the dataset according to the procedure employed by Breitling *et al*.^54^, Bailey *et al*.^55^, Graham *et al*.^6,10^, and Garayburu-Caruso *et al*.^42^. The pairwise mass differences between each detected metabolite were determined and matched to a database of 1298 frequently observed biochemical transformations (**Supplemental File 4**). For example, if the mass difference between two metabolites was 18.0343, that would putatively indicate a loss or gain of an ammonium group, while a mass difference of 103.0092 would putatively indicate loss or gain of a cysteine. This calculation is enabled by the ultrahigh mass resolution of FTICR-MS data; given this resolution, we considered any between-metabolite mass difference within 1 ppm of the expected mass of a transformation to be a match. This analysis provides two outputs: a transformation profile outlining the number of times a putative transformation could occur in a given sample and pairwise mass difference between every peak. Multivariate similarities between the transformation profiles of each sample were visualized by generating a Jaccard dissimilarity-based non-metric multidimensional scaling (NMDS) ordination (*metaMDS*, ‘vegan’ package v2.5-6)^56^. Using these pairwise mass differences and transformation associations, we then generated a transformation network in which nodes are metabolites and edges are transformations (**Supplemental Figure 1**)^11,57,58^. Relationships between metabolites were determined by first selecting the largest cluster of interconnected nodes (discarding everything not within this cluster) and measuring the stepwise distance between each pair of metabolites (i.e., the minimum number of transformations required to connect one metabolite to another metabolite within the largest cluster of the biochemical transformation network). These pairwise distances were then standardized between 0 and 1.

Relationships among metabolites were also evaluated using a number of metabolite characteristics estimated from inferred molecular formulae. To do so elemental composition (C-, H-, O-, N-, S-, P-content), double-bond equivalents (DBE), modified aromaticity index (AI_mod_), and Kendrick’s defect were used as metabolite characteristics indicating molecular composition and structure of the metabolites. These metrics were combined to generate a pairwise Euclidean distance matrix with each distance representing approximate dissimilarity (i.e., further distances indicate less similar metabolites). These molecular differences were then weighted by the previously measured transformation distances that were themselves scaled to be between 0 and 1. A UPGMA hierarchical clustering analysis was then used to convert this combined distance matrix into a dendrogram which approximates the relationships among metabolites (**Supplemental File 5**). This resulted in the transformation weighted molecular characteristics dendrogram (TWCD). While Danczak *et al*.^11^ used three different dendrograms, doing so is beyond the scope of the current study and we chose to use the TWCD as it integrates more information relative to other dendrogram methods.

### Diversity Analyses

The metabolite data were treated as an assemblage of ecological units following the methodology outlined in Danczak *et al*.^11^. All metabolites were treated on a presence/absence basis – peak intensities were not used due to charge competition^47,52^. Richness measurements and Jaccard-based dissimilarity metrics (*vegdist*, ‘vegan’ package 2.5-6)^56^ were used to assess the compositional differences among metabolite assemblages. The TWCD was used to measure dendrogram-based alpha-diversity indices including Faith’s PD (*pd*, ‘picante’ package v1.8)^59^, mean nearest taxon distance (MNTD), mean pairwise distance (MPD), variance in nearest taxon distance (VNTD), and variance in pairwise distance (VPD) (*generic*.*metrics*, ‘pez’ package v1.2-0)^60–65^. β-mean nearest taxon distance (βMNTD) was measured using the *comdistnt* function in the picante R package^59^. Jaccard dissimilarity and βMNTD results were visualized using a principal coordinates analysis (PCoA; *pcoa*, ‘ape’ package v5.3)^66^.

### Ecological Null Modeling

Null modeling was performed to quantify the relative influences of variable selection, homogeneous selection, and stochastic processes over metabolite assemblages^11^. Specifically, the β-Nearest Taxon Index (βNTI) was calculated to measure the influence of stochastic and deterministic assembly processes^12,13,15^. βNTI was estimated for each pairwise assemblage comparison. To do so, a null distribution of 999 βMNTD values were generated and compared to the observed βMTND value for a given pair of assemblages. Pairwise comparisons with |βNTI| > 2 indicate that deterministic processes were responsible for observed differences in metabolite composition. In contrast, pairwise comparisons with |βNTI| < 2 indicate that stochastic processes were responsible for observed differences in metabolite composition.

Furthermore, the deterministic processes can be separated into two classes. When βNTI > 2, differences in metabolite composition are greater than would be expected by random chance (i.e., greater than the stochastic expectation). This is analogous to ‘variable selection,’ which occurs when deterministic processes drive divergence in composition between a pair of assemblages^13,14^. When βNTI < 2, differences in metabolite composition are less than the stochastic expectation. This is analogous to ‘homogeneous selection,’ which occurs when deterministic process drive convergence in composition between a pair of assemblages. Mean βNTI values for each sample were obtained and used in all analyses and plots.

### Statistics and Plot Generation

Differences in distributions (i.e., diversity analyses, molecular properties) were evaluated using Mann Whitney U tests (*wilcox*.*test*, ‘stats’ package). Multivariate differences (i.e., ordinations) were identified using PERMANOVA tests (*adonis*, vegan package v2.5-6)^56^. All correlations were Spearman-based and were performed using the *rcorr* function (‘Hmisc’ package v4.2)^67^. All boxplots and scatter/line plots were generated using the ‘ggplot2’ R package (v3.2.1)^68^; three-dimensional ordinations were generated using the ‘plot3D’ R package (v1.1.1)^69^.

All R scripts used within this manuscript are available on GitHub at https://github.com/danczakre/HJA-FTICR-Ecology. The uncalibrated, peak-picked FTICR-MS files and aqueous geochemistry data are available at https://data.ess-dive.lbl.gov/view/doi:10.15485/1509695^70^. The FTICR-MS report used in this study has been included as **Supplemental Data 1**.

## Acknowledgments

Pacific Northwest National Laboratory is operated by Battelle Memorial Institute for the U.S. Department of Energy under Contract No. DE-AC05-76RL01830. This research was supported by the U.S. Department of Energy (DOE), Office of Biological and Environmental Research (BER), as part of BER’s Subsurface Biogeochemistry Research Program (SBR). This contribution originates from the SBR Scientific Focus Area (SFA) at the Pacific Northwest National Laboratory (PNNL). A portion of this work was performed at the Environmental Molecular Science Laboratory User Facility.

## Author Contributions

R.E.D, A.E.G., E.B.G, and J.C.S. conceptualized the study. V.A.G., J.W.M., L.R., and J.R.W. collected samples and analyzed anions/cations. R.K.C, J.G.T., and N.K collected FTICR-MS data and assisted with analyses. S.P.H. and A.S.W. assisted with hydrological interpretations. R.E.D. performed the ecological and statistical analyses. R.E.D. drafted the manuscript but all authors contributed to the writing

## Supplemental Legend

Supplemental Figure 1: Visual representation of the transformation network utilized to generate the transformation-weighted characteristics dendrogram (TWCD). Each node within the network represents an individual metabolite while the edges connecting each node is a transformation. Note the large cluster of interconnected nodes near the middle of the plot.

Supplemental File 1: Metadata and geochemistry for the field site at Watershed 1 (WS1) in the HJ Andrews Experimental Forest.

Supplemental File 2: Significant Spearman-based correlations between average sample βNTI and site geochemistry. The table is short given that only significant correlations are provided. Correlations labeled “Bulk” indicate that both surface water and pore water samples were considered in correlations (i.e., the entire dataset), correlations labeled “SW48” were performed only with surface water samples, and correlations labeled “PP48” were performed only using pore water samples.

Supplemental File 3: Significant correlations between average sample βNTI and putative biochemical transformations. Sheet 1 includes those significant correlations between individual transformation relative proportions and βNTI, while Sheet 2 are all correlations between transformation groups and βNTI (i.e., not only significant correlations). As in Supplemental File 2, correlations labeled “Bulk” indicate that both surface water and pore water samples were considered in correlations (i.e., the entire dataset), correlations labeled “SW48” were performed only with surface water samples, and correlations labeled “PP48” were performed only using pore water samples.

Supplemental File 4: Database of known and frequently observed biochemical transformations. This file is used to identify putative biochemical transformations using ultrahigh-resolution mass differences obtained from FTICR-MS datasets.

Supplemental File 5: The transformation-weighted characteristics dendrogram (TWCD) obtained using the UPGMA hierarchical clustering method.

Supplemental Data 1: Aligned and calibrated FTICR-MS report generated using Formularity.

