## Supplementary figures and images for "Ecological theory applied to environmental metabolomes reveals compositional divergence despite conserved molecular properties"

### Supplemental Figure 1

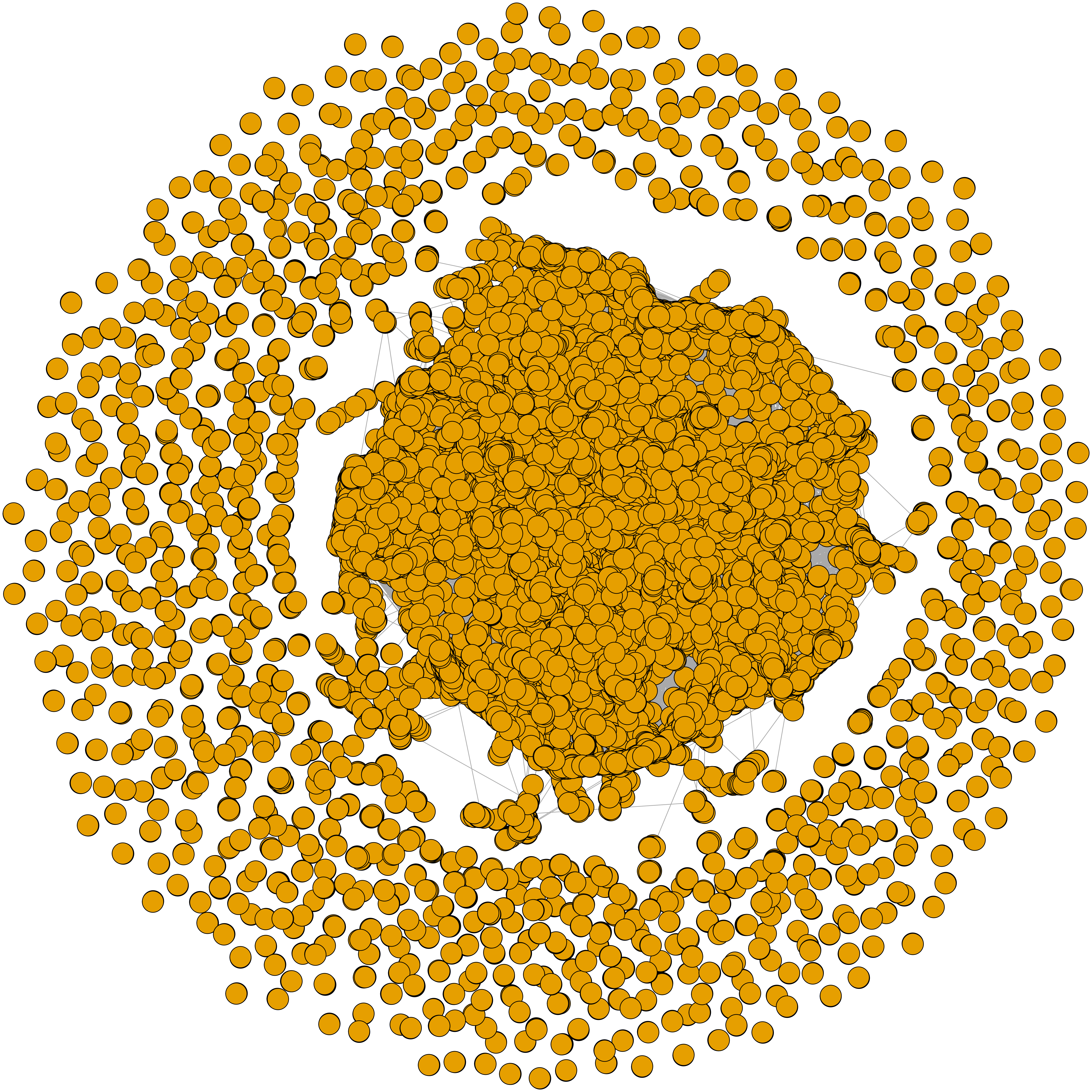
